# Social information used to elicit cache protection differs between pinyon jays and Clark’s nutcrackers

**DOI:** 10.1101/2021.02.28.433225

**Authors:** Alizée Vernouillet, Dawson Clary, Debbie M. Kelly

## Abstract

Behavioral plasticity can be described as the ability to adjust behavior depending on environmental information. We used a food-storing (caching) paradigm, during which individuals either ate or cached food under different conditions, to investigate whether they could adjust their caching behavior when observed by conspecifics and heterospecifics, and which cues they used to elicit these behavioral changes. We examined the location and number of caches made by two corvid species differing in sociality, highly social pinyon jays (*Gymnorhinus cyanocephalus*) and less social Clark’s nutcrackers (*Nucifraga columbiana*). Although pinyon jays cached a similar amount of food across conditions, they allocated more caches to areas less accessible to the observer. Nutcrackers, however, reduced the number of seeds cached when another nutcracker was present in comparison to when they cached alone. Both species relied on different social cues to elicit re-caching: pinyon jays responded to the amount of time the observer spent close to the caching locations, whereas nutcrackers responded to the amount of time the observer spent pilfering their caches. The differences in cache protection behaviors and the social cues eliciting them may be explained by the species’ social organization. Pinyon jays may only adjust their caching behavior when necessary, as they are often surrounded by other individuals. Clark’s nutcrackers reduce their caching when observed, as they have more opportunities to cache alone, and may resort to additional cache protection when experiencing pilferage. Overall, our results provide insight into understanding how pressures associated with the social environment may influence foraging behaviors.

## Introduction

Behavioral plasticity can be defined as the ability to adjust behavioral responses to cope with physical (both biotic and abiotic) and social changes in the environment (Mery & Burns, 2009). Individuals display a range of behaviors in response to different environmental conditions. Adaptive behaviors are then favored by evolutionary processes and overtime, these suites of behaviors spread within a species (Lande 2009, Chevin & Lande 2010). Hence, behavioral plasticity varies within and between species. More specifically, the degree of behavioral plasticity displayed by a species may depend on the complexity and unpredictability of the environment, with more varied environments selecting for greater behavioral plasticity (reviewed by Komers 1997; Snell-Rood & Ehlman 2021).

Foraging strategies, for instance, can often demonstrate behavioral plasticity (Morand-Ferron & Giraldeau 2010). Caching, the behavior of storing food when resources are abundant for later consumption when resources become scarce (Vander Wall 1990), is an example of such behavioral plasticity, as it requires adapting to an unstable food source. For this behavior to be evolutionary advantageous, caching individuals must retrieve a sufficient proportion of the food they previously stored (Krebs 1990). Thus, caching individuals must not only relocate their caches but also protect these resources from potential thieves (*pilferers*) (Dally *et al*. 2006). Indeed, many corvid (e.g., crows, magpies, jays) and parid (e.g., chickadees) species have excellent observational spatial memory and can retain this information to relocate caches made by others, sometimes days after a cache has been made (e.g., pinyon jays *Gymnorhinus cyanocephalus*, Bednekoff & Balda 1996a; Mexican jays *Aphelocoma wollweberi* and Clark’s nutcrackers *Nucifraga columbiana*, Bednekoff & Balda 1996b; ravens *Corvus corax*, Scheid & Bugnyar 2008; great tits *Parus major*, Brodin & Urhan 2014). Thus, to reduce the likelihood of their caches being pilfered, caching individuals may engage in a suite of *cache protection behaviors*, during which caching behavior is adjusted depending on the perceived pilfering risk. As such, investigations focusing on caching and cache protection behaviors may be informative for the study of behavioral plasticity.

Cache protection behaviors can involve a modification of the amount of cached food or a preference of where food is cached. Examples of modification of the amount of food cached include: reduced caching to prevent food loss (e.g., California scrub jays *Aphelocoma californica*, Dally *et al*. 2005; Clark’s nutcrackers, Clary & Kelly 2011; Canada jays *Perisoreus canadensis*, Martin & Sherry 2021), and enhanced caching to compensate for food loss (e.g., Eurasian jays *Garrulus glandarius*, Bossema 1979). Examples of spatial preference for the allocation of cached food include: caching out-of-view of observing birds (e.g., ravens, Bugnyar & Kotrschal 2002; Clark’s nutcrackers, Tornick & Gibson 2021), allocating caches to less risky locations (e.g., pinyon jays, Vernouillet *et al*. 2021a), re-caching to a different location after the observing bird has left (e.g., California scrub jays, Emery *et al*. 2004; Clark’s nutcrackers Clary & Kelly 2011), and using the properties of the substrate (e.g., color, acoustics) to reduce cues available to observing conspecifics (e.g., California scrub jays, Stulp *et al*. 2009; Kelley & Clayton 2017). Which cache protection behaviors are used and when to display them showcase behavioral plasticity in a changing social environment, as a uniform response to all observers, regardless of the threat they pose, may be costly for the individual.

The aforementioned studies examining cache pilfering by avian species have mainly investigated the interactions among conspecifics, with limited investigations of pilfering by heterospecifics (but see Pravosudov 2008). However, heterospecific pilfering has been reported in the wild on multiple occasions (e.g., Burnell & Tomback 1985; Dally *et al*. 2006; Chock *et al*. 2019), and may account for a major proportion of caches lost to pilfering (e.g., Dittel *et al*. 2017; Penner & Devenport 2011; Swift *et al*. 2021; Vander Wall & Jenkins 2003). Additionally, learning to adjust caching behavior in the presence of a heterospecific observer might showcase the behavioral plasticity of caching individuals. Indeed, caching individuals need to first determine a heterospecific observer as a pilfering threat and then flexibly adjust their caching behavior according to the behavioral tendencies of individuals of that species.

Corvids are a taxonomic family of birds that have adapted to disparate environments, and are renowned for their acts of behavioral plasticity (reviewed in Emery & Clayton 2004; Taylor 2014; Lambert *et al*. 2018). Reliance on cached food varies widely amongst corvid species, from non-caching white-throated magpie-jay (*Calocitta formosa*) to long-term specialized cachers such as pinyon jays and Clark’s nutcrackers (de Kort *et al*. 2006). The propensity to cache seems to be innate in most corvid species as it is hypothesized that the common ancestor of all corvids was a moderate cacher (i.e., caching different food items through the year but not being entirely dependent upon those caches; de Kort & Clayton 2006). However, more advanced caching behaviors that improve caching efficiency, such as cache protection behaviors, may require learning (Emery & Clayton 2001; Dally *et al*. 2006). Yet, whether corvids can determine if a heterospecific observer is a potential pilfering threat, and engage in behavioral plasticity through cache protection behaviors when observed by a heterospecific, remains unknown.

To date, most studies examining cache protection strategies by corvids have focused on moderately social species (e.g., California scrub jays, Emery *et al*. 2004, Dally *et al*. 2005) and less social species (e.g., nutcrackers Clary & Kelly 2011; Tornick & Gibson 2021; ravens, Bugnyar & Kotrschal 2002; Canada jays, Martin & Sherry 2021; Eurasian jays, Shaw & Clayton 2012). In contrast, few studies have addressed the challenges encountered by highly social species (but see, pinyon jays: Bednekoff & Balda 1996a; Vernouillet *et al*. 2021a), and no studies have directly compared the caching behavior and the social cues used to elicit changes in caching behavior between highly social and less social species using the same paradigm. Here, we examined the caching behavior of highly-social pinyon jays and of less-social Clark’s nutcrackers when in the presence of a conspecific observer, as well as when in the presence of a heterospecific observer.

Pinyon jays and Clark’s nutcrackers are two closely-related (Ericson *et al*. 2005) corvid species that have similar foraging ecologies. Their distributions overlap in the southwestern United States of America, where they inhabit woodlands (Marzluff & Balda 1992; Tomback 1998). Both pinyon jays and Clark’s nutcrackers rely heavily on thousands of pine seeds cached during winter and early spring (around 26,000 caches and 33,000 caches, respectively; Vander Wall & Balda 1981; Vander Wall 1990). Since these species are sympatric, instances of individual nutcrackers briefly joining flocks of pinyon jays during late summer and early fall (i.e., when they collect pine seeds to cache) have previously been observed (Balda *et al*. 1972). Hence, heterospecific pilfering of caches between species during that period is possible.

Although similar in their foraging ecology, both species differ in their social structures. Pinyon jays are a highly-social species, living in large flocks of up to hundreds of individuals with high fission-fusion dynamics (Marzluff & Balda 1992; Clayton & Emery 2008), whereas Clark’s nutcrackers are a less-social species, whose social group typically consists of the mating pair and the offspring of the year (Clayton & Emery 2008; Tomback 1998). The differing social structures of the two species likely influences the social pressures under which they cache. For instance, pinyon jays are known to make caching trips in groups (Marzluff & Balda 1992), and as such, are usually surrounded by conspecifics when caching. Clark’s nutcrackers, as a less-social species, prefer to eat and make their caches away from other individuals (Tomback 1998). Hence, each species may have evolved different cache protection behaviors according to the demands of their social structure and may differ in the cues that elicit cache protection behaviors.

During this study, we aimed to address three research objectives. First, we evaluated which cache protection behaviors pinyon jays and Clark’s nutcrackers use when faced with a pilfering threat. With the specter of a replication crisis in the field of animal cognition (Farrar *et al*. 2020), even in caching studies (e.g., Amodio *et al*. 2021a, b), this step is an important first check onto validating our results. Based on previous research, we predicted that when observed caching, pinyon jays would allocate their caches to safer places (Vernouillet *et al*. 2021a), whereas Clark’s nutcrackers would reduce their caching (Clary & Kelly 2011). Second, we evaluated whether pinyon jays and Clark’s nutcrackers would respond to a heterospecific observer as a potential cache threat. Given that both species are highly dependent on cached food, we expected both species to adjust their caching behavior in the presence of a heterospecific observer, to lower the pilfering risk to their caches. Third, we examined which cues affected the caching behavior of pinyon jays and Clark’s nutcrackers, and if these cues differed between species considering their respective social structures. We expected that less-social Clark’s nutcrackers may rely on the presence/absence of an observer as their natural caching ecology may accustom them to wait for opportunities to cache while alone. Highly-social pinyon jays, however, might use more subtle behavioral cues, as they are afforded less opportunities for caching in privacy in natural settings.

## Methods

### Subjects

Pinyon jays and Clark’s nutcrackers served as caching birds (herein referred as “cachers”; *n* = 10 for each species, five female and five male pinyon jays and six female and four male Clark’s nutcrackers) or as observing birds (herein referred as “observers”; *n* = 4 two females for each species). All individuals were captured as adults from populations around Flagstaff (Arizona, USA) and were in captivity for approximately seven to ten years prior to this study. Five of the nutcracker cachers had previous experience with caching paradigms (Clary & Kelly 2011, 2016a, b); the five other nutcracker cachers, and all ten pinyon jay cachers had previous unrelated experimental experience (e.g., concept learning (Vernouillet *et al*. 2021b; Wright *et al*. 2017; exploratory behaviour, Vernouillet & Kelly 2020).

All birds were housed in individual cages for the duration of the study (pinyon jays: 51 x 51 x 72 cm, nutcrackers: 82 x 54 x 76 cm, width x depth x height, respectively), with multiple wood perches, at the University of Manitoba, Canada. Pinyon jays were housed in a colony room alongside California scrub jays, whereas nutcrackers were kept in a species-specific colony room. All colony rooms were maintained at 22°C with a 12:12 day-night cycle, with light onset at 0700 (Central Daylight time). These temperature and lighting conditions correspond to the average temperature and photoperiod during September in Flagstaff (USA), the peak caching season for pinyon jays and Clark’s nutcrackers. Birds were given *ad libitum* water and grit. When not experiencing the food restriction procedure (see General procedures below), birds were fed a regular diet consisting of a mixture of parrot pellets, turkey starter, sunflower seeds, mealworms, peanuts, powder of oyster shells, and the vitamin powder supplement Prime®. Birds were monitored and weighed daily to ensure a healthy weight during the experiment.

All applicable international, national, and institutional guidelines for the care and use of animals were followed. Our research protocol was approved by University of Manitoba’s Animal Care Committee (#F2014-037) and complied with the guidelines set by the Canadian Council on Animal Care.

### Caching Apparatus

The caching apparatus was the same as used in Vernouillet *et al*. (2021a). The experiment was conducted in an experimental room, separated from the colony room. Birds were individually tested in a cage (123.5 x 63.5 x 74.5 cm, width x depth x height), divided into two equally-sized compartments. The entire cage was surrounded by white curtains (Figure S1; also Figure 1 from Vernouillet *et al*. 2021a). A transparent acrylic divider separated the two compartments, each of which contained a perch. One compartment served as the “caching compartment”, whereas the other served as the “observing compartment”.

**Figure 1.**
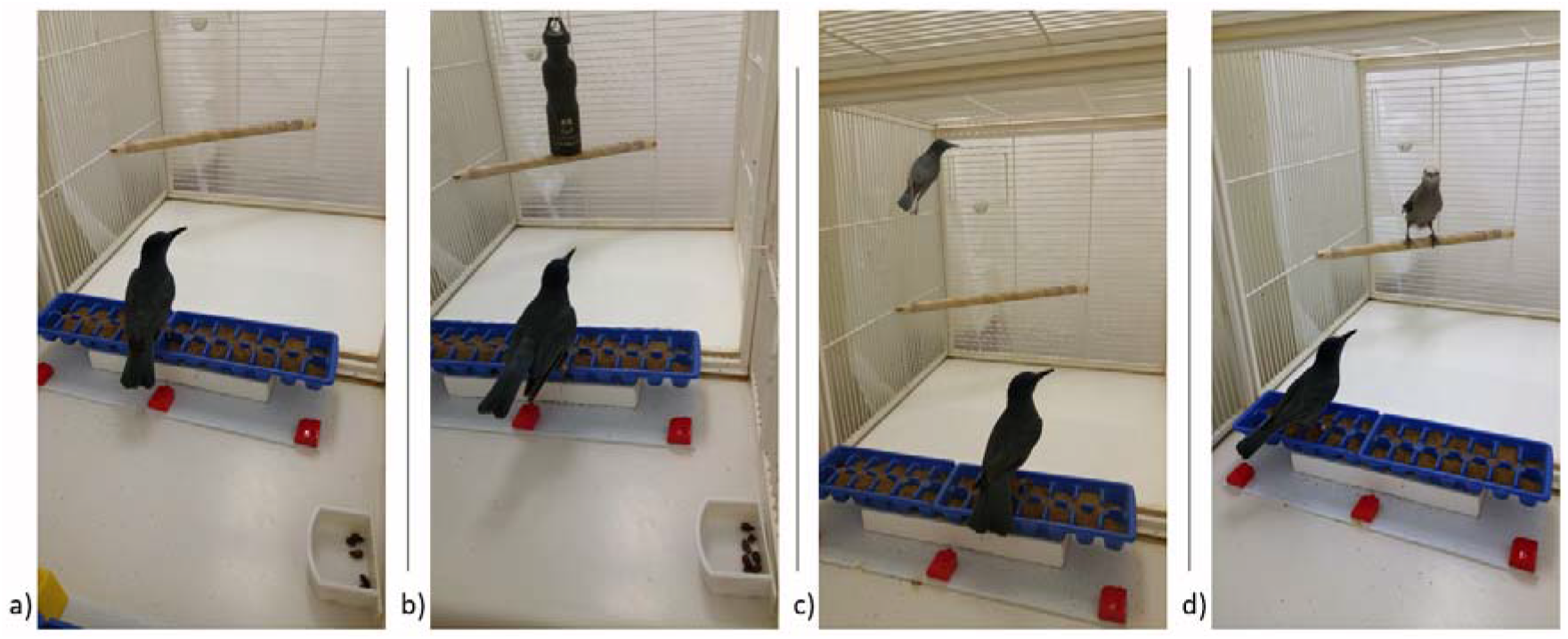
Pictorial representation of the experimental conditions during the *Caching phase*. The caching compartment contained the cacher (nearest bird, in this example, a pinyon jay) and the observing compartment either: **a)** remained empty (*Alone* condition), **b)** contained an object (*Object* condition), **c)** contained a conspecific (in this example, another pinyon jay; *Conspecific* condition), or **d)** contained a heterospecific (in this example, a Clark’s nutcracker; *Heterospecific* condition).

Within the caching compartment, cachers were given two plastic ice cube trays (49.5 x 11 cm, length x width) comprised of 2 rows, each with 13 wells, filled with sand. Trays were made visually distinctive by affixing coloured plastic MegaBlocks™ at the base of each tray. Each cacher received the same pair of distinct trays throughout the study, but the arrangement of colored blocks differed across subjects. One tray (hereafter referred to as the “Pilfered Tray”) was positioned parallel and flush against the acrylic cage divider, whereas the other tray (hereafter referred to as the “Safe Tray”) was positioned on the opposite wall of the caching compartment (i.e., parallel to the first tray). The arrangement of colored MegaBlocks for each tray remained consistent throughout the experiment, permitting the birds to reliably differentiate between the two trays. A food dish was placed in the caching compartment beside the cage door, between the two trays (Figure S1). All trials were recorded using an EverFocus® 1/3” color digital camera positioned either beside or above the experimental cage and using the motion tracking program, BiObserve® through Windows XP.

### General Procedures

The general experimental procedures were similar to Vernouillet *et al*. (2021a). Briefly, each bird was given a weekly experimental session. Cachers were food deprived for 24h before the start of the *Caching phase*, by removing all the food from their home cage. This procedure created food uncertainty and motivated individuals to cache (Clary & Kelly 2011; Kamil & Balda 1985; Vernouillet *et al*. 2021a). The session started with a 45-min *Caching phase*, during which the cacher was provided with a dish of 50 pine nuts to cache or consume. The observing compartment was arranged as per the condition to be completed that session (see Conditions below). This phase was immediately followed by a 3-min *Pilfering phase*, during which the Pilfered Tray was placed in the observing compartment, alongside the divider and in the same orientation as during the *Caching phase*. The Safe Tray was placed on a stool outside of both compartments (remaining visible, but inaccessible to both birds). Upon the completion of the *Pilfering phase*, all birds were returned to the colony room. After a one-hour delay, the cacher was given a 5-min *Re-caching phase*, during which both trays were returned to the cacher, such that they could eat or re-cache (i.e., move a pine seed to a different location from where it was previously cached) some of the seeds while alone. A re-cache was thus recorded when a pine seed was found in a different well of either the same tray or the opposite tray, compared to where it was originally placed. Upon completion of the *Re-caching phase*, the cacher was returned to the colony room and provided with a small amount of food to ensure a healthy bodyweight was maintained, while ensuring the bird would remain motivated to retrieve its caches the following day. During the *Retrieving phase*, which occurred 24 hours after the completion of the *Caching phase*, the cacher was placed in the caching compartment with the safe and Pilfered Trays, unaltered from the previous *Re-caching phase*. The cacher was permitted 45 minutes to consume or re-cache the previously cached pine seeds, during which the observing compartment of the caching cage was empty (i.e., retrieval was always conducted while alone). Additional *Retrieving phases* were administered, on the following day, every three hours, if necessary, until the cacher recovered the entirety of its caches. After each phase, the researcher recorded the number and location of each pine seed (for more details, see Supplementary Material, or Vernouillet *et al*. 2021a).

#### Conditions

Each cacher first experienced (at least) three weeks of baseline trials during which individuals experienced the above procedures without an observer present (similar to the *Alone* condition). These baseline trials were used to establish whether individuals cached reliably and whether they were distributing their cached seeds equally both trays (i.e., no tray preference; see additional methodology details in the Supplementary Material). After the baseline trials, each cacher experienced three blocks of four conditions (*Alone*, *Conspecific, Heterospecific*, and *Object*), with each condition randomly selected and not repeated within the same block. Each condition refers to what (if anything) was present in the observing compartment during the *Caching* and *Pilfering phases*. Observers did not participate as cachers to exclude potential effects of pilfering experience on their caching behaviour (Emery & Clayton 2001). The pairing of observers to cacher remained consistent throughout the experiment (each observer watched five cachers of each species).

##### Alone

During the *Caching phase*, the observing compartment remained empty (Figure 1a). During the *Pilfering phase*, the Pilfered Tray was placed in the empty observing compartment for three minutes, and the pine seeds cached in that tray remained undisturbed. This condition was conducted to assess the baseline caching behaviour of cachers and was used to assess whether exposure to the other experimental conditions changed the caching behaviour of individuals while alone.

##### Object

During the *Caching phase*, an inanimate object (a black water bottle, 27.5 x 7.0 cm, height x diameter) was affixed to the perch in the observing compartment (Figure 1b). During the *Pilfering phase*, the Pilfered Tray was placed in the observing compartment with the object for three minutes, whereas the Safe Tray was placed on a stool outside of the cage, remaining visible to the cacher. The experimenter removed pine seeds from the Pilfered Tray after the *Pilfering phase* (but before the *Re-caching phase*). Amount of pilfering was kept consistent across individuals by removing 33% of pine seeds cached during the first and second blocks, and 50% of pine seeds cached during the third block, to resemble natural variation of pilfering in the wild (pilfer amounts were chosen according to Vander Wall & Jenkins 2003). Both trays were given back to the cacher during the *Re-caching phase*. This condition served as a control to determine whether the cacher modified its caching behaviour in response to cache loss, and when compared with the *Conspecific* and *Heterospecific* conditions, assessed the importance of social and motion cues in the display of cache protection strategies.

##### Conspecific

During the *Caching phase*, a bird of the same species as the cacher occupied the observing compartment (Figure 1c). During the *Pilfering phase*, the Pilfered Tray was placed in the observing compartment with the observer, who was given three minutes to access the tray in view of the cacher. The Safe Tray was placed on an inaccessible stool outside of the cage, but visible to both birds. If the observer did not pilfer enough seeds according to the pilfering criterion described for the Object condition, additional pine seeds were removed by the experimenter after the *Pilfering phase* (but before the *Re-caching phase*). Similarly, if the observer pilfered more seeds than the standardized threshold, the experimenter added caches back in the tray after the *Pilfering phase* (but before the *Recaching phase*). For both cases, the experimenter adjusted the seed number such that all areas of the tray were pilfered equally. This condition was conducted to assess whether the presence and behaviour of a conspecific observer influenced the cacher’s behaviour.

##### Heterospecific

This condition was conducted as during the *Conspecific* condition, with the exception that the bird in the observing compartment was of a different species than the cacher (Figure 1d). This condition was conducted to assess whether the presence and the behaviour of a heterospecific observer influenced the cacher’s behaviour.

### Behavioural Measures

#### Dependent variables

We evaluated the potential cache protection behaviours (i.e., consumption, enhanced/reduced caching, preferential allocation of caches, and re-caching; Table 1) used by pinyon jays and Clark’s nutcrackers when observed by conspecifics and by heterospecifics. During the *Caching phase*, we examined the overall number of pine seeds cached to evaluate for enhanced/reduced caching, the number of pine seeds cached in the trays to evaluate for preferential allocation, and the number of pine seeds eaten to evaluate for consumption. During the *Re-caching phase*, we examined the proportion of re-cached seeds to evaluate for re-caching, which corresponded to the proportion of pine seeds found in new locations (compared to the locations after the *Caching phase*) out of all the remaining pine seeds at the end of the *Re-caching phase*. All measures were evaluated at a global-level (i.e., with respect to the total number of pine seeds cached) and at a tray-level, to determine whether cachers associated the pilfering risk with the Pilfered Tray, but not the Safe Tray. At the tray level, the proportion of re-cached pine seeds corresponded to the proportion of pine seeds that were found in new locations out of all the seeds remaining in each tray at the end of the *Re-caching phase* (compared to the locations after the *Caching Phase*).

**Table 1.** Cache protection behaviors that were evaluated for pinyon jays and Clark’s nutcrackers when caching alone, in presence of an object, or when observed by a conspecific or a heterospecific. The factor Block and its interactions (Block x Condition, Block x Condition x Tray) accounted for the possibility of individuals learning the risk associated with each condition over the course of the study. For all models, the identity of the cacher was included as a random factor.

| Cache protection behaviours (adapted from Dally *et al.*, 2006) | Description | Predicted significant factors in models |
| --- | --- | --- |
| Consumption | Increase eating pine seeds when observed | Eaten pine seeds ~ Condition/Time Interacting<br>Eaten pine seeds ~ Condition x Time Interacting |
| Enhanced/reduced caching | Increase/decrease caching when observed | Cached pine seeds ~ Condition/Time Interacting<br>Cached pine seeds ~ Condition x Time Interacting |
| Preferential allocation | Increase/decrease caching in high-risk locations | Cached pine seeds ~ Tray<br>Cached pine seeds ~ Tray x Condition/Time Interacting |
| Re-caching | Move high-risk pine seeds to new sites | Re-caching ~ Condition/Time Interacting/Time Pilfering<br>Re-caching ~ Condition x Time Interacting/Time Pilfering<br>Re-caching ~ Tray<br>Re-caching ~ Tray x Condition/Time Interacting/Time Pilfering |

#### Behaviour of the observer

We assessed whether cachers would modify their caching behaviour in the mere presence of an observer or whether the behaviour of the observer when caching influenced the caching behaviour of the birds. Hence, we measured the duration (in seconds) the observer spent interacting with the cacher during the *Caching phase* and with the Pilfered Tray during the *Pilfering phase* from the recorded sessions of the *Conspecific* and *Heterospecific* conditions. During the *Caching phase*, the observer was defined as interacting with the cacher when the observer was standing within the closest third section of the observing compartment nearest the divider separating the two compartments, but could not access any caching trays. During the *Pilfering phase*, the observer was defined as interacting with the Pilfered Tray when searching in the tray, retrieving pine seeds, or standing on the tray.

### Statistical Analyses

We performed analyses separately for each species. We used a generalized linear mixed model (GLMM) approach for our analyses, with the number of seeds cached and eaten during the *Caching phase* and the proportion of seeds re-cached during the *Re-caching phase* as dependent variables. Condition, duration of time the observer spent interacting with the cacher during the *Caching phase*, and duration of time the observer spent interacting with the Pilfered Tray during the *Pilfering phase* were included as fixed factors in our models to assess whether the cacher responded to the observer’s presence and/or behaviour when caching during the *Caching* and *Re-caching phases*. Another fixed factor included in the models was experimental block to detect any evidence of behavioural change in caching over time. We also performed separate analyses on the number of seeds cached in each tray and the proportion of seeds re-cached in each tray to determine whether the cachers treated the two trays differently. For these analyses, we included tray identity (pilfered vs. safe) as a fixed factor. Identity of the cacher was included in all models as a random factor to account for repeated measures taken on each cacher. Residual plots indicated assumptions of normality were met for the number of seeds cached and eaten during the *Caching phase*. For the *Recaching phase*, we used a binomial approach to determine which factors affected the probability that seeds were re-cached. Analyses were conducted using R version 3.3.2 (R Core Team 2013) with the *lme4* (Bates *et al*. 2015), *lsmeans* (Lenth 2016) and *Rmisc* (Hope 2013) packages.

To assess whether specific factors influenced our dependent variables, we compared full models to a reduced model that did not include the factor that was evaluated (Harrison *et al*. 2018; Tredennick *et al*. 2021). Fixed factors included in the models were tray, condition, block, amount of time the observer interacted with the cacher during the *Caching phase*, amount of time the observer interacted with the Pilfered Tray during the *Pilfered phase* (only for the analyses on re-cached seeds), as well as the interactions block x condition, condition x tray, condition x block x tray, condition x amount of time the observer interacted with the cacher during the *Caching phase*, condition x amount of time the observer interacted with the Pilfered Tray during the *Pilfered phase* (only for the analyses on re-cached seeds), tray x amount of time the observer interacted with the cacher during the *Caching phase*, tray x amount of time the observer interacted with the Pilfered Tray during the *Pilfered phase*. Parameter estimation was achieved using residual maximum likelihood ratio test. Post-hoc analyses were conducted using pairwise comparisons. *P*-values of post-hoc analyses were adjusted using the Tukey method to consider multiple comparisons (Lenth 2016). Alpha was set at 0.05 for all statistical analyses.

## Results

### Pinyon Jays

#### Enhanced/reduced caching

There was no statistical evidence that pinyon jays engaged in enhanced or reduced caching when observed, as there were no differences in the overall number of pine seeds cached by pinyon jays between conditions (*M ± SE: Alone:* 14.6 ± 2.4, *Object:* 15.0 ± 2.1, *Conspecific:* 15.6 ± 2.2, *Heterospecific:* 16.2 ± 2.2 pine seeds cached; *χ^2^_(3)_* = 0.143, *p* = 0.986; Figure 2; Table S1). No other fixed factors examined in our analyses explained the total number of seeds cached during the *Caching phase* by the pinyon jays (Table S1).

**Figure 2.**
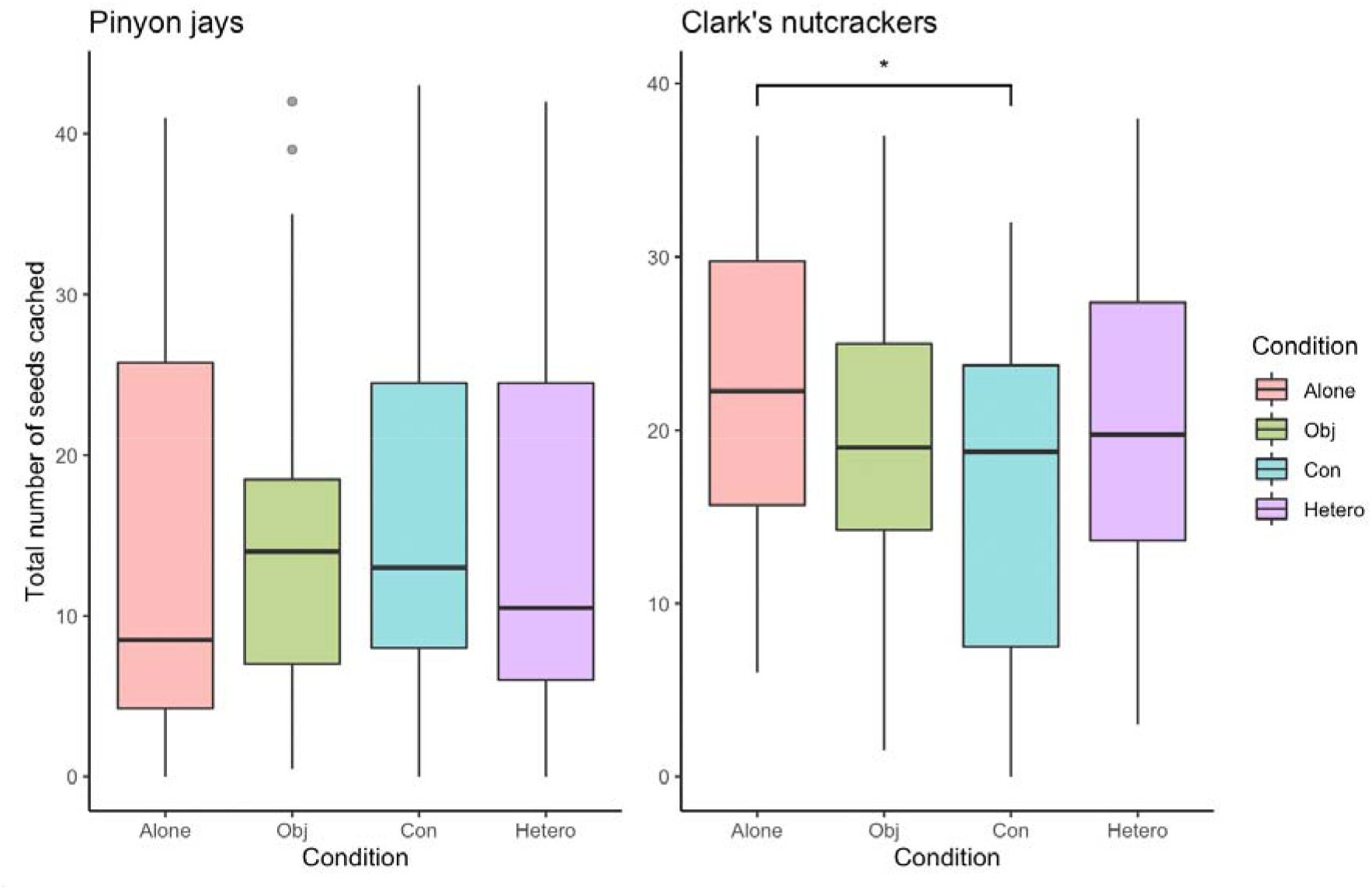
Total number of pine seeds cached by pinyon jays (left) and Clark’s nutcrackers (right) during the *Caching phase* during the *Alone, Object (Obj), Conspecific (Con)*, and *Heterospecific (Hetero*) conditions. There was no difference in the number of seeds cached by pinyon jays across conditions (*p* = 0.986). Nutcrackers cached less when in the presence of another nutcracker (*i.e., Conspecific* condition) in comparison to when they were caching alone (*p* = 0.005).

#### Preferential allocation of caches

We found that pinyon jays preferentially allocated their caches in the Safe Tray (*M ± SE*: Pilfered Tray: 5.7 ± 0.5 pine seeds, Safe Tray: 9.6 ± 0.8 pine seeds; *χ^2^_(1)_* = 24.489, *p* < 0.001; Figure S2). However, they did not preferentially allocate their caches depending on the condition *χ^2^_(6)_* = 2.187, *p* = 0.902; Figure 3), blocks (*χ^2^_(4)_* = 4.574, *p* = 0.334; Figure S3), nor was there a significant condition x block interaction (*χ^2^_(12)_* = 8.625, *p* = 0.735). No other fixed factors examined in our analyses explained the number of seeds cached per tray during the *Caching phase* by the pinyon jays (Table S1).

**Figure 3.**
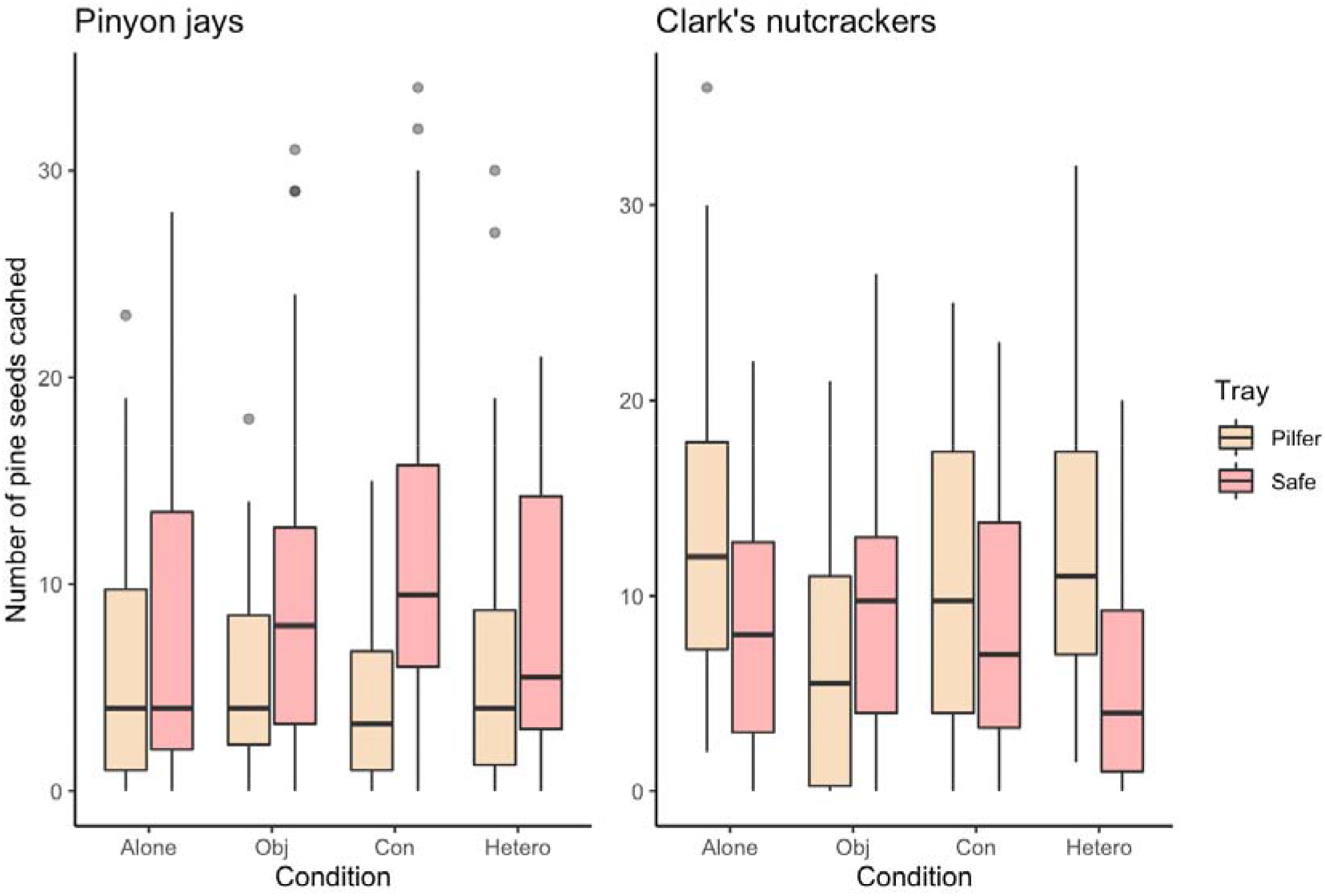
Number of pine seeds cached in the Pilfered Tray and in the Safe Tray by pinyon jays (left) and Clark’s nutcrackers (right) during the *Caching phase*. There was no statistical evidence that the number of pine seeds cached differed between trays depending on the condition in pinyon jays (*p* = 0.902) nor in Clark’s nutcrackers (*p* = 0.128).

#### Consumption

There was no statistical evidence that pinyon jays engaged in consumption as a cache protection behaviour during our study (see Supplementary Material, Table S2, Figure 4).

**Figure 4.**
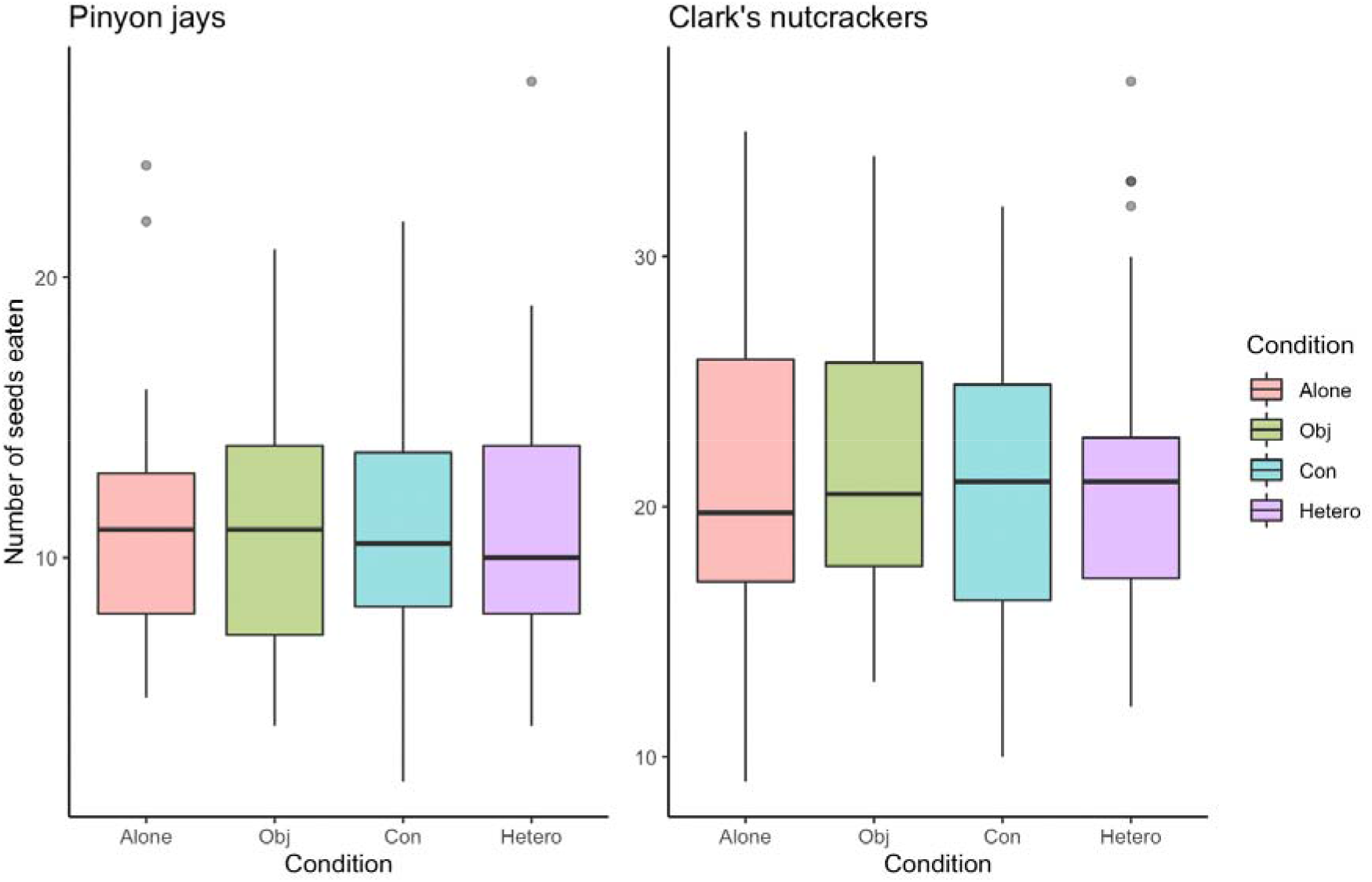
Number of pine seeds eaten by pinyon jays (left) and by Clark’s nutcrackers (right) during the *Caching phase*. There was no difference in the number of seeds eaten by pinyon jays (*p* = 0.884) and by Clark’s nutcrackers (*p* = 0.855) across conditions.

#### Overall re-caching

We found evidence that pinyon jays used re-caching as a cache protection behaviour. The overall proportion of re-cached pine seeds during the *Re-caching phase* by pinyon jays was significantly influenced by the interaction between the amount of time the observer spent interacting with the cacher during the *Caching phase* and condition *χ^2^_(1)_* = 12.313, *p* < 0.001; Table S3). When observed (i.e., during the *Conspecific* and *Heterospecific* conditions), pinyon jay cachers increased the overall proportion of re-cached pine seeds when the pinyon jay observers and nutcracker observers had spent more time interacting with the trays during the *Caching* phase (Pinyon jay observer estimate: 0.03 ± 0.01/min; Clark’s nutcracker observer estimate: 0.01 ± 0.01/min; Figure S6).

The overall proportion of re-cached pine seeds during the *Re-caching phase* also differed in certain trials, as the interaction between block and condition was significant in our models *χ^2^_(6)_* = 24.099, *p* < 0.001; Table S3). Post-hoc analyses indicated that this result was mostly driven by one trial, with a significantly lower proportion of re-cached seeds during the second block of the *Object* condition than any other condition or block (see Supplementary Materials for more details). No other fixed factors examined in our analyses explained the overall proportion of re-cached pine seeds during the *Re-caching phase* by the pinyon jays (Table S3).

#### Preferential re-caching

The proportion of pine seeds re-cached by pinyon jays differed in each tray, with a greater proportion of caches identified in new locations of the Pilfered Tray in comparison to the Safe Tray (*M ± SE*: Pilfered Tray: 0.45 ± 0.04, Safe Tray: 0.23 ± 0.03 *χ^2^_(1)_* = 64.496, *p* < 0.001; Table S3; Figure 5). Yet, the proportion of re-cached pine seeds in each tray did not differ among conditions (*χ^2^_(6)_* = 2.980, *p* = 0.811; Figure S5). However, when accounting for the effects of block, the proportion of cached seeds within the tray that were found in new locations was higher in the Pilfered Tray in comparison to the Safe Tray *χ^2^_(12)_* = 31.034, *p* = 0.002), more specifically during the *Heterospecific* condition across all blocks (*z* > 3.730, *p* < 0.002 for all comparisons), during the *Conspecific* condition during the last block (*z* = 4.541, *p* < 0.001), and during the *Object* condition during the last block (*z* = 3.449, *p* < 0.001). No other fixed factors examined in our analyses explained the proportion of re-cached pine seeds per tray during the *Re-caching phase* by the pinyon jays (Table S3).

**Figure 5.**
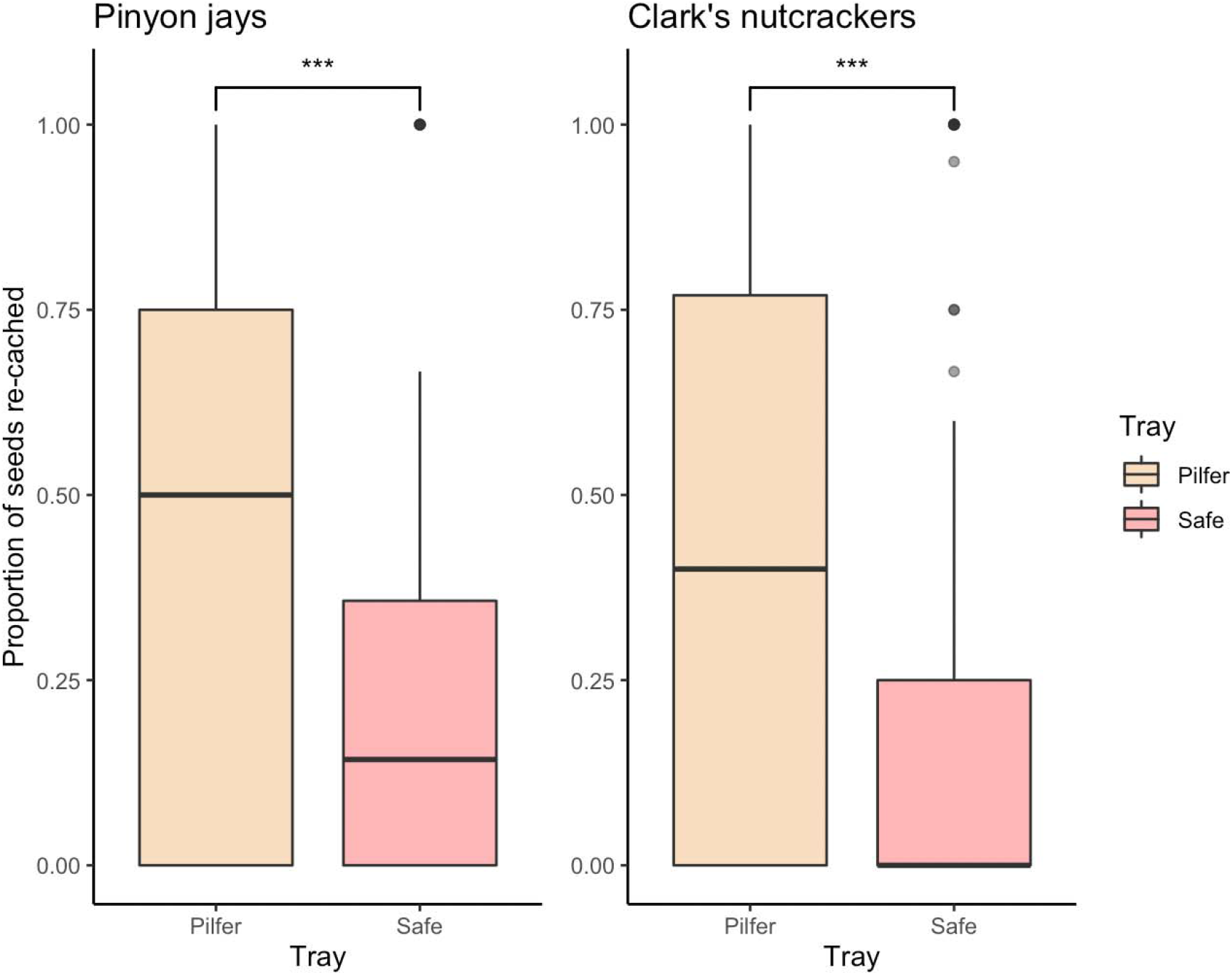
Proportion of re-cached pine seeds per tray by pinyon jays (left) and by Clark’s nutcrackers (right) during the *Re-caching phase*. Both pinyon jays and Clark’s nutcrackers had a higher proportion of re-cached seeds in the Pilfered Tray than in the Safe Tray (*p* < 0.001 for both species).

### Clark’s Nutcrackers

#### Enhanced/reduced caching

We found some statistical evidence that Clark’s nutcrackers modify the number of caches made based on whether they were observed during caching. The overall number of pine seeds cached by nutcrackers was best explained by condition *χ^2^_(3)_*) = 7.916, *p* = 0.048; Table S4; Figure 2). Nutcrackers cached significantly fewer pine seeds when observed by a conspecific compared to when caching alone (*M ± SE: Alone:* 22.0 ± 1.6; *Conspecific:* 16.2 ± 1.8 pine seeds; *t* = 3.39, *p* = 0.005), and tended to cache fewer pine seeds when observed by a conspecific than when observed by a heterospecific (i.e., pinyon jay), though this difference was not significant (*M ± SE*: *Heterospecific:* 20.3 ± 1.8 pine seeds; *t* = −2.37, *p* = 0.089). There was no difference in the number of pine seeds cached when caching alone and when observed by a heterospecific (*t* = 1.02, *p* = 0.741). No other fixed factors examined in our analyses explained the overall number of pine seeds during the *Caching phase* by the Clark’s nutcrackers (Table S4).

#### Preferential allocation of caches

We found that Clark’s nutcrackers preferentially allocated their caches. However, unlike pinyon jays, nutcrackers cached a higher number of pine seeds in the Pilfered Tray than in the Safe Tray, as there was a main effect of Tray (*M ± SE:* Safe Tray: 8.4 ± 0.6 pine seeds, Pilfered Tray: 10.9 ± 0.7 pine seeds; *χ^2^_(1)_* = 8.212, *p* = 0.004; Figure S2), regardless of condition (*χ^2^_(6)_* = 9.918, *p* = 0.128; Figure 3). There was also a main effect of block on the number of pine seeds cached in each tray by the nutcrackers (Table S4). Nutcrackers cached more pine seeds in the Pilfered Tray than in the Safe Tray during the third (i.e., final) block (*M ± SE*: Pilfered Tray: 12.9 ± 1.4 pine seeds, Safe Tray: 6.8 ± 0.8 pine seeds; *t* = 4.198, *p* < 0.001), but not during the first block (*M ± SE*: Pilfered Tray: 10.7 ± 1.1 pine seeds, Safe Tray: 9.4 ± 1.0 pine seeds; *t* = 0.922, *p* = 0.358) nor the second block (*M ± SE*: Pilfered Tray: 9.0 ± 1.2, Safe Tray: 9.0 ± 1.2 pine seeds; *t* = 0.047, *p* = 0.963). No other fixed factors examined in our analyses explained the number of seeds cached per tray during the *Caching phase* by the Clark’s nutcrackers (Table S4).

#### Consumption

There was no statistical evidence that Clark’s nutcrackers engaged in consumption as a cache protection behaviour during our study (see Supplementary Material, Table S5, Figure 4).

#### Overall re-caching

We found evidence that Clark’s nutcrackers used re-caching as a cache protection behaviour. When observed (i.e., during the *Conspecific* and the *Heterospecific* conditions), Clark’s nutcracker cachers increased the overall proportion of recached pine seeds as nutcracker observers and pinyon jay observers spent more time pilfering the Pilfered Tray during the *Pilfering* phase (Overall estimate: 0.11 ± 0.06/min; *χ^2^_(1)_* = 7.062, *p* = 0.008; Figure S5; Table S6). The proportion of re-cached pine seeds also depended on the species of the observer that was pilfering, with nutcracker cachers significantly increasing the proportion of pine seeds re-cached as conspecific observers spent more time pilfering the pilfered tray, but this was not the case with heterospecific observers (Clark’s nutcracker observer estimate: 0.14 ± 0.08/min; Pinyon jay observer estimate: 0.07 ± 0.09/min; *χ^2^_(1)_* = 7.942, *p* = 0.005; Table S6).

The overall proportion of re-cached seeds also differed across conditions and blocks, as the interaction between blocks and conditions was significant in our models (*χ^2^_(6)_* = 31.175, *p* < 0.001; Table S6). Proportion of re-cached seeds was significantly higher during the second block of the *Object* condition, in comparison to the second block of the *Alone* condition (*z* = −3.013, *p* = 0.014) and the *Heterospecific* condition (*z* = −3.796, *p* < 0.001), and significantly higher during the third block of the *Heterospecific* condition in comparison to the *Alone* condition (*z* = −2.898, *p* = 0.020). No other fixed factors examined in our analyses explained the overall proportion of re-cached pine seeds during the *Re-caching phase* by the Clark’s nutcrackers (Table S6).

#### Preferential re-caching

The proportion of re-cached pine seeds by nutcrackers differed in each tray, with a greater proportion of re-caches identified in new locations of the Pilfered Tray in comparison to the Safe Tray (*M ± SE*: Pilfered Tray: 0.43 ± 0.04, Safe Tray: 0.19 ± 0.03; *χ^2^_(1)_* = 64.496, *p* < 0.001; Figure 5; Table S6). Similar to the overall proportion of re-cached pine seeds, when observed (i.e., during the *Conspecific* and the *Heterospecific* conditions), an increased proportion of re-cached pine seeds were found in new locations of the Pilfered Tray and a decreased proportion of re-cached pine seeds were found in new locations of the Safe Tray as nutcracker and pinyon jay observers spent more time pilfering during the *Pilfering* phase (Pilfered Tray estimate: 0.56 ± 0.09/min, Safe Tray estimate: 0.39 ± 0.10/min; *χ^2^_(2)_* = 7.051, *p* = 0.029; Table S6).

Additionally, nutcrackers re-cached a significantly greater proportion in the Pilfered Tray than in the Safe Tray for every condition and across blocks (*z* > 2.730, *p* < 0.006 for all comparisons), except for the second and third blocks of the *Object* condition (*z* = 1.380, *p* = 0.168 and *z* = 1.823, *p* = 0.068 for the second and third blocks, respectively). No other fixed factors examined in our analyses explained the proportion of re-cached pine seeds per tray during the *Re-caching phase* by the Clark’s nutcrackers (Table S7).

## Discussion

Our study evaluated three main research objectives. Firstly, to replicate prior findings, we investigated which cache protection behaviors were used by highly-social pinyon jays and less-social Clark’s nutcrackers. Overall, the cache protection behaviors we observed for both species are consistent with those found during previous studies. Pinyon jays cached similar number of seeds regardless of observer (see also Vernouillet *et al*. 2021a), whereas nutcrackers reduced the number of seeds cached in the presence of a conspecific (see also Clary & Kelly 2011). Additionally, pinyon jays placed their caches in safe locations throughout the study, whereas nutcrackers showed no statistical preference between trays, except for a preference to cache in the ‘risky’ pilfered tray during the final block of trials. For both pinyon jays and Clark’s nutcrackers, compared to the safe tray, a greater proportion of the seeds present in the pilfered tray after the *Re-caching phase* were re-caches. Hence, both species behaved as if the caches in the pilfered tray were at a greater risk of being pilfered. Yet, each species may have assessed the risk associated with each tray differently, with pinyon jays showing greater aversion than nutcrackers to caching in the pilfered tray.

Secondly, we evaluated whether each species would engage in cache protection behaviors in the presence of a heterospecific. The pinyon jays may have been more affected by the presence of a heterospecific, as evidenced by their preferential re-caching in the pilfered tray throughout the study, whereas this preference was only seen during the final block of trials when in the presence of a conspecific. Clark’s nutcrackers, however, only suppressed caching in response to conspecific, but not heterospecific observers. One potential explanation for this strong response to the presence of observing nutcrackers for both species may be that observing nutcrackers were perceived as more of a pilfering threat than pinyon jay observers, despite experiencing the same standardized cache loss. Indeed, observing nutcrackers spent more time close to the caching trays during the *Caching phase* than observing pinyon jays (Table S7).

The potential difference in threat level posed by each species may also relate to species differences in body size and competition in the wild. Nutcrackers tend to be larger than pinyon jays (average weight range for Clark’s nutcrackers: 106-160g, for pinyon jays: 90-120g; Tomback 1998; Johnson & Balda 2020), so nutcrackers may not have perceived pinyon jays as a threat to their caches. The two species also differ in their likelihood of interactions with heterospecifics in the wild, as nutcrackers tend to inhabit higher elevations than pinyon jays, in a harsher, and potentially less competitive, habitat (Tomback 1998, Marzluff & Balda 1992).

Thirdly, we evaluated which cues each species use to modify their cache protection behavior. The tendency for birds to re-cache depended on different social cues from the observer for both species. For pinyon jays, the proportion of re-caches depended on the duration of time the observer spent close to the cacher during the caching session. For nutcrackers, the proportion of re-caches depended on the duration of time the observing bird spent pilfering the pilfered tray during the pilfering phase. As such, both species demonstrated behavioral plasticity by adjusting their cache protection behaviors based on the social information available (i.e., the behavior of the observer towards the cacher).

Social information can be exchanged between conspecifics and heterospecifics when multiple species share the same territory or resources. For example, some species are influenced by heterospecifics when responding to alarm calls (Dutour & Danel 2020; Magrath *et al*. 2017), when choosing a breeding site (Morinay *et al*. 2020; Tolvanen *et al*. 2020), or even when foraging (Romero-Gonzalez *et al*. 2020). For our study species, individual Clark’s nutcrackers briefly join flocks of pinyon jays during late summer and early fall, when both species collect pine seeds to cache for the winter (Balda *et al*. 1972). Both species’ caches are susceptible to pilfering by one another, but also by Steller’s jays (*Cyanocitta stelleri)*, another social jay species (Burnell & Tomback 1985). Both species interact with each other for a limited amount of time, as nutcrackers spend most of the year at higher elevations than jays (Marzluff & Balda 1992; Tomback 1998). However, no aggressive interactions between wild pinyon jays and nutcrackers have been published. Interactions between the two species would likely be concentrated during the caching period. Here, we provide experimental evidence that both corvid species can use the social information provided by an observer, regardless of the species, and adjust their caching behavior according to the observer’s behavior.

To date, only one other study evaluated whether a caching species, the mountain chickadee (*Poecile gambeli*), can adjust its caching behavior in the presence of a pilfering heterospecific, the red-breasted nuthatch (*Sitta canadensis*; Pravosudov 2008). Mountain chickadees cached preferentially in hidden sites in the presence of observing nuthatches but used both hidden and visible sites when they were alone, suggesting chickadees interpreted nuthatches as a threat to their caches. Likewise, our results support the idea that species that rely heavily on cached items can adjust their caching behavior using social information provided by both conspecifics and heterospecifics.

As with cache protection behaviors (Vernouillet *et al*. 2021a), the social cues that elicit them may align with the demands of social living. Since highly social pinyon jays rarely cache in private (Marzluff & Balda 1992), it may be more advantageous for an individual to adjust its behavior only when warranted by the behavior of the observer, and not simply in the presence of an observer. Therefore, pinyon jays may use nuanced social information during caching to modify their behavior, due to the more restrictive conditions in which they cache, regardless of whether the observer pilfers the caches afterwards (Vernouillet *et al*. 2021a). On the contrary, less social species such as Clark’s nutcrackers and Eurasian jays (*Garrulus glandarius*) can reduce their caching in the presence of another bird (Clary & Kelly 2011; Shaw & Clayton 2012), as they have more opportunities to cache in private, and adjust their re-caching behavior only once experiencing pilferage. This difference in the degree of reliance on personal and social information is supported by a recent study comparing two corvid species that differed in sociality (McCune *et al*. 2022). Social Mexican scrub jays relied more on social information to solve a motor task, whereas relatively less social California scrub jays relied more on personal information. Similarly, pinyon jays were faster at learning a motor task in a social setting than by themselves, whereas nutcrackers performed equally well in both settings (Templeton *et al*. 1999). Our results also support the idea that both highly social and less social species use social information to adjust their caching behavior, but less social species may rely more on their personal experience of being pilfered to elicit more involved forms of cache protection behaviors, such as re-caching. Which specific behaviour of the observer that caching birds attend to (e.g., distance from caching locations, gaze direction, direct interactions with the cacher, dominance displays), perhaps dependent on sociality, warrants further investigation.

In summary, we investigated the caching behavior of highly social pinyon jays and less social Clark’s nutcrackers. When observed by a conspecific, highly social pinyon jays preferentially allocated cached food in areas less accessible to the observer, and less social Clark’s nutcrackers reduced the number of caches made. Both species also increased recaching, especially for caches that were at greater risk of pilferage. These cache protection behaviors can be linked to the social ecology of each species and reflect the social environment during caching. When observed by a heterospecific, pinyon jays displayed the same cache protection behavior as when they are observed by a conspecific, whereas Clark’s nutcrackers only increased re-caching in response to pilfering. This result might reflect that nutcrackers have less opportunities to be pilfered by a heterospecific in the wild, but may also suggest that observing nutcrackers were more threatening than observing pinyon jays due to the size difference between species. Finally, when evaluating which cues elicited cache protection behavior in both species, we found that cachers from both species adjusted their caching behavior based on the behavior of the observer. Pinyon jays responded to the amount of time the observer spent near a caching tray, whereas Clark’s nutcrackers responded to the amount of time the observer spent pilfering their caches. Once again, this difference in cues used aligns with the social environment individuals experience during caching. Pinyon jays may only adjust their caching behavior when necessary, as signaled by the behavior of the observer during caching, as they are often surrounded by other individuals. Clark’s nutcrackers reduce their caching when observed and resort to additional cache protection behaviors when experiencing pilferage. Together, these results suggest that corvids may differ in their use of social information and in the extent to which they interpret heterospecific observers as a threat. These findings may indicate that the ability to view a conspecific and a heterospecific as a threat is shared among corvids regardless of social organization. Our results provide insight into how ecological pressures associated with the social environment and caching may influence the behavioral plasticity of corvids.

