## Supplementary Tables and Figures for "Social information used to elicit cache protection differs between pinyon jays and Clark’s nutcrackers"

Alizée Vernouillet is now a Postdoctoral Researcher at the Department of Experimental Psychology at Ghent University.

AV and DMK developed the study; AV conducted the experiments and analysed the data; AV wrote the manuscript with comments from DC and DMK.

Research support was provided by a University of Manitoba Graduate Fellowship and by a BOF postdoc fellowship (#BOF.PDO.2021.0035.01) to AV, and by a Natural Science and Engineering Research Council Discovery grant (#RGPIN/4944-2017) and Canada Research Chair to DMK. The authors are grateful to Iroshini Gunasekara, Thomas Rawliuk, and Nicole Tongol for helping with video scoring, and to Ben Farrar and two anonymous reviewers for their comments. Data are available as supplementary material. The experiment was not preregistered.

**Methods**

*Experimental Procedures*

Prior to the start of the experiment, individuals were given at least three caching sessions to determine whether individuals cached reliably (as determined by caching at least 5 pine seeds per session for three consecutive sessions). This baseline session was also used to determine if the individual had a bias towards caching within a specific tray. Potential bias towards one tray was assessed by calculating whether the proportion of seeds was significantly different between trays using a binomial test, with one tray being favored over the other for three consecutive sessions. None of the birds showed a bias towards one tray and none of the birds needed more than three baseline trials. After these three baseline trials, the birds proceeded to the experiment the following week.

During the *Caching* *phase*, the cacher was provided with a dish of 50 pine seeds to cache or to consume within a 45-minute period. The cacher was also given two plastic ice cube trays (49.5 x 11 cm, length x width) each with 2 rows of 13 wells filled with sand. Trays were made visually distinctive by affixing coloured plastic MegaBlocks™ at the base of each tray. The observing compartment was arranged as per the condition to be completed that session (see Conditions in the main text): it would either be empty (*Alone* condition), contain a bottle (*Object* condition), contain a conspecific observer (*Conspecific* condition), or contain a heterospecific observer (*Heterospecific* condition). At the end of the *Caching phase*, the two trays were removed from the caching compartment to allow the researcher to record the number and location of each pine seed, while the cacher remained in the caching compartment.

A three-minute *Pilfering* *phase* occurred immediately after the pine seeds were counted. During the *Pilfering* *phase*, the Pilfered Tray was placed in the observing compartment, alongside the divider and in the same orientation as during the *Caching phase*. This tray was accessible to the observer present in the observing compartment during the *Conspecific* and the *Heterospecific* conditions. The Safe Tray was placed on a stool outside of both compartments (remaining visible, but inaccessible to both birds). Upon the completion of the *Pilfering phase*, both birds were returned to the colony room.

One hour later, the cacher experienced a *Re‑caching* *phase* of five minutes in the caching compartment, during which both trays were returned to the cacher, such that they could eat or re-cache some of the seeds while alone. The *Re-caching* *phase* was instituted to provide the birds with an opportunity to change the location of their seeds or to retrieve them after witnessing the pilfering event, while remaining satiated from the prior *Caching phase*. This allowed us to differentiate the birds' cache protection behaviours from behaviours motivated by hunger. Upon completion of the *Re‑caching* *phase*, the cacher was returned to the colony room and provided with a small amount of food to support the maintenance of a healthy bodyweight, while ensuring the bird would still be motivated to retrieve its caches the following day.

During the *Retrieving* *phase*, which occurred 24 hours after the completion of the *Caching* *phase*, the cacher was placed in the caching compartment with the Safe and Pilfered Trays, unaltered from the previous *Re-caching phase*. The cacher was permitted 45 minutes to consume or re‑cache the previously cached pine seeds, during which the observing compartment of the caching cage was empty (i.e., retrieval was always conducted while alone). Additional *Retrieving phases* were administered, on the following day, every three hours if necessary, until the cacher recovered the entirety of its caches. At the end of each *Retrieving phase*, the number and location of the pine seeds cached were noted, together with the re-caching occurrences.

**Results**

*Consumption of seeds*

There was no statistical evidence that pinyon jays engaged in consumption as a cache protection behaviour during our study, as there were no statistical differences in the number of pine seeds eaten across conditions (*χ^2^_(3)_* = 0.654; *p* = 0.884; Figure 4; Table S2). No other fixed factors examined in our analyses explained the number of seeds eaten during the *Caching phase* by the pinyon jays (Table S2).

There was no statistical evidence that Clark’s nutcrackers engaged in consumption as a cache protection behaviour during our study, as there were no statistical differences in the number of pine seeds eaten across conditions during the *Caching* phase (*χ^2^_(3)_* = 0.776, *p* = 0.855; Figure 4; Table S5). No other fixed factors examined in our analyses explained the number of seeds eaten during the *Caching phase* by the Clark’s nutcrackers (Table S5).

*Overall re-caching*

The overall proportion of re-cached pine seeds during the *Re-caching phase* also differed in certain trials, as the interaction between block and condition was significant in our models (*χ^2^_(6)_* = 24.099, *p* < 0.001; Table S3). Post-hoc analyses indicated that this result was mostly driven by one trial, with a significantly lower proportion of re-cached seeds during the second block of the *Object* condition (see Supplementary Materials), both in comparison to the other conditions (*z* > 3.115, *p* < 0.010, for all comparisons), and compared to the first and third blocks of the *Object condition* (z > 3.290, *p* < 0.003, for all comparisons). No other trials significantly differed from one another. No other fixed factors examined in our analyses explained the overall proportion of re-cached pine seeds during the *Re-caching phase* by the pinyon jays (Table S3).

*Behaviour of the Observers*

During the *Caching phase,* pinyon jay observers spent significantly less time interacting with pinyon jay cachers than nutcracker observers did (Table S7). During the *Pilfering phase*, there was no significant difference in the duration of time spent interacting with the pilfered tray between pinyon jay and nutcracker observers (Table S8).

During the *Caching phase,* pinyon jay observers also spent significantly less time interacting with nutcracker cachers compared to nutcracker observers (Table S7). During the *Pilfering phase*, there was also no significant difference in the duration of time pinyon jay and nutcracker observers spent interacting with the pilfered tray (Table S7).

**Table S1.** Factors evaluated to explain the number of pine seeds cached by pinyon jays during the *Caching phase*. Significant factors are indicated in bold.

|  | *χ^2^* | *d.f.* | *p* |
| --- | --- | --- | --- |
| Condition | 0.143 | 3 | 0.986 |
| Block | 1.041 | 2 | 0.594 |
| **Tray** | **24.489** | **1** | **< 0.001** |
| Time Interacting | 0.225 | 1 | 0.635 |
| Condition x Block | 3.426 | 6 | 0.754 |
| Condition x Time Interacting | 0.216 | 1 | 0.642 |
| Tray x Condition | 2.187 | 2 | 0.511 |
| Tray x Block | 4.575 | 4 | 0.334 |
| Tray x Condition x Block | 5.456 | 12 | 0.941 |
| Tray x Time Interacting | 1.341 | 2 | 0.511 |

**Table S2.** Factors evaluated to explain the number of pine seeds eaten by pinyon jays during the *Caching phase*.

|  | *χ^2^* | *d.f.* | *p* |
| --- | --- | --- | --- |
| Condition | 0.654 | 3 | 0.884 |
| Block | 0.512 | 2 | 0.774 |
| Time Interacting | 1.004 | 1 | 0.316 |
| Condition x Block | 10.169 | 6 | 0.118 |
| Condition x Time Interacting | 0.845 | 1 | 0.358 |

**Table S3.** Factors evaluated to explain the proportion of pine seeds re-cached by pinyon jays during the *Re-Caching phase*.

|  | *χ^2^* | *d.f.* | *p* |
| --- | --- | --- | --- |
| Condition | 1.847 | 3 | 0.605 |
| Block | 1.749 | 2 | 0.417 |
| **Tray** | **20.008** | **1** | **< 0.001** |
| Time Interacting | 0.000 | 1 | 1.000 |
| Time Pilfering | 0.591 | 1 | 0.442 |
| **Condition x Block** | **24.099** | **6** | **< 0.001** |
| **Condition x Time Interacting** | **12.313** | **1** | **< 0.001** |
| Condition x Time Pilfering | 3.354 | 1 | 0.067 |
| Tray x Condition | 2.980 | 6 | 0.811 |
| Tray x Block | 6.683 | 4 | 0.154 |
| **Tray x Condition x Block** | **31.034** | **12** | **0.002** |
| Tray x Time Interacting | 1.000 | 1 | 0.317 |
| Tray x Time Pilfering | 4.089 | 2 | 0.130 |

**Table S4.** Factors evaluated to explain the number of pine seeds cached by Clark’s nutcrackers during the *Caching phase*.

|  | *χ^2^* | *d.f.* | *p* |
| --- | --- | --- | --- |
| **Condition** | **7.916** | **3** | **0.048** |
| Block | 2.117 | 2 | 0.347 |
| **Tray** | **8.212** | **1** | **0.004** |
| Time Interacting | 0.000 | 1 | 0.998 |
| Condition x Block | 4.582 | 6 | 0.599 |
| Condition x Time Interacting | 0.992 | 1 | 0.319 |
| Tray x Condition | 9.918 | 6 | 0.128 |
| **Tray x Block** | **13.301** | **4** | **0.010** |
| Tray x Condition x Block | 8.625 | 12 | 0.735 |
| Tray x Time Interacting | 0.331 | 2 | 0.847 |

**Table S5.** Factors evaluated to explain the number of pine seeds eaten by Clark’s nutcrackers during the *Caching phase*.

|  | *χ^2^* | *d.f.* | *p* |
| --- | --- | --- | --- |
| Condition | 0.776 | 3 | 0.855 |
| Block | 0.850 | 2 | 0.654 |
| Time Interacting | 0.023 | 1 | 0.881 |
| Condition x Block | 4.701 | 6 | 0.583 |
| Condition x Time Interacting | 0.147 | 1 | 0.701 |

**Table S6.** Factors evaluated to explain the proportion of pine seeds re-cached by Clark’s nutcrackers during the *Re-Caching phase*.

|  | *χ^2^* | *d.f.* | *p* |
| --- | --- | --- | --- |
| Condition | 7.780 | 3 | 0.051 |
| Block | 3.875 | 2 | 0.144 |
| **Tray** | **64.496** | **1** | **< 0.001** |
| Time Interacting | 1.942 | 1 | 0.163 |
| **Time Pilfering** | **7.062** | **1** | **0.008** |
| **Condition x Block** | **31.175** | **6** | **< 0.001** |
| Condition x Time Interacting | 1.636 | 1 | 0.201 |
| **Condition x Time Pilfering** | **7.942** | **1** | **0.005** |
| Tray x Condition | 11.179 | 6 | 0.083 |
| Tray x Block | 5.382 | 4 | 0.250 |
| **Tray x Condition x Block** | **33.736** | **12** | **< 0.001** |
| Tray x Time Interacting | 1.700 | 1 | 0.192 |
| **Tray x Time Pilfering** | **7.051** | **2** | **0.029** |

**Table S7.** Average duration in minutes (±*SE*) pinyon jay and Clark’s nutcracker observers spent interacting with the cacher during the *Caching phase* and with the pilfered tray during the *Pilfering phase*. Range of durations are presented between brackets.

| Cacher | Phase | Observers | | *t* | *p* |
| --- | --- | --- | --- | --- | --- |
|  |  | Pinyon jay | Nutcracker |  |  |
| Pinyon jay | Caching | 4.53 ± 1.12 (0.00 – 23.15) | 17.36 ± 1.85 (0.00 – 37.26) | -5.939 | < 0.001 |
|  | Pilfering | 1.26 ± 0.25 (0.00 – 3.13) | 1.64 ± 0.27 (0.00 – 3.38) | 1.300 | 0.099 |
| Nutcracker | Caching | 3.09 ± 1.04 (0.00 – 21.7) | 17.84 ± 1.95 (2.13 – 39.45) | -6.686 | < 0.001 |
|  | Pilfering | 1.21 ± 0.24 (0.00 – 3.10) | 1.18 ± 0.22 (0.00 – 3.43) | 0.253 | 0.401 |

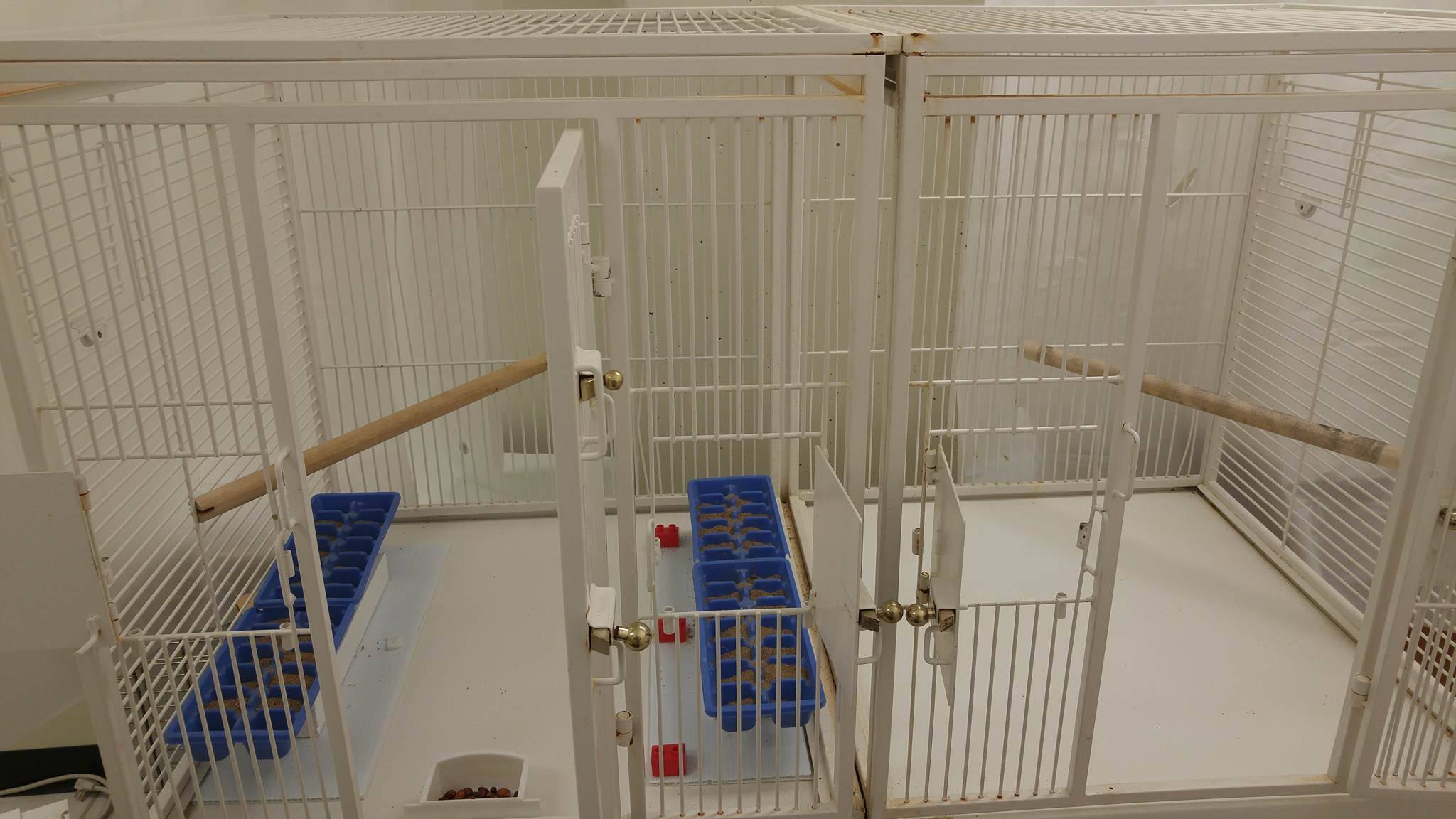

**Figure S1.** The caching apparatus used in the experiment. The caching bird was placed in the left compartment (*caching compartment*), whereas the adjoining compartment (*observing compartment*) was either empty (*Alone* condition, as shown here), contained a black plastic bottle secured to the perch (*Object* condition), an observing pinyon jay or an observing Clark’s nutcracker. The two compartments were separated by a transparent acrylic divider.

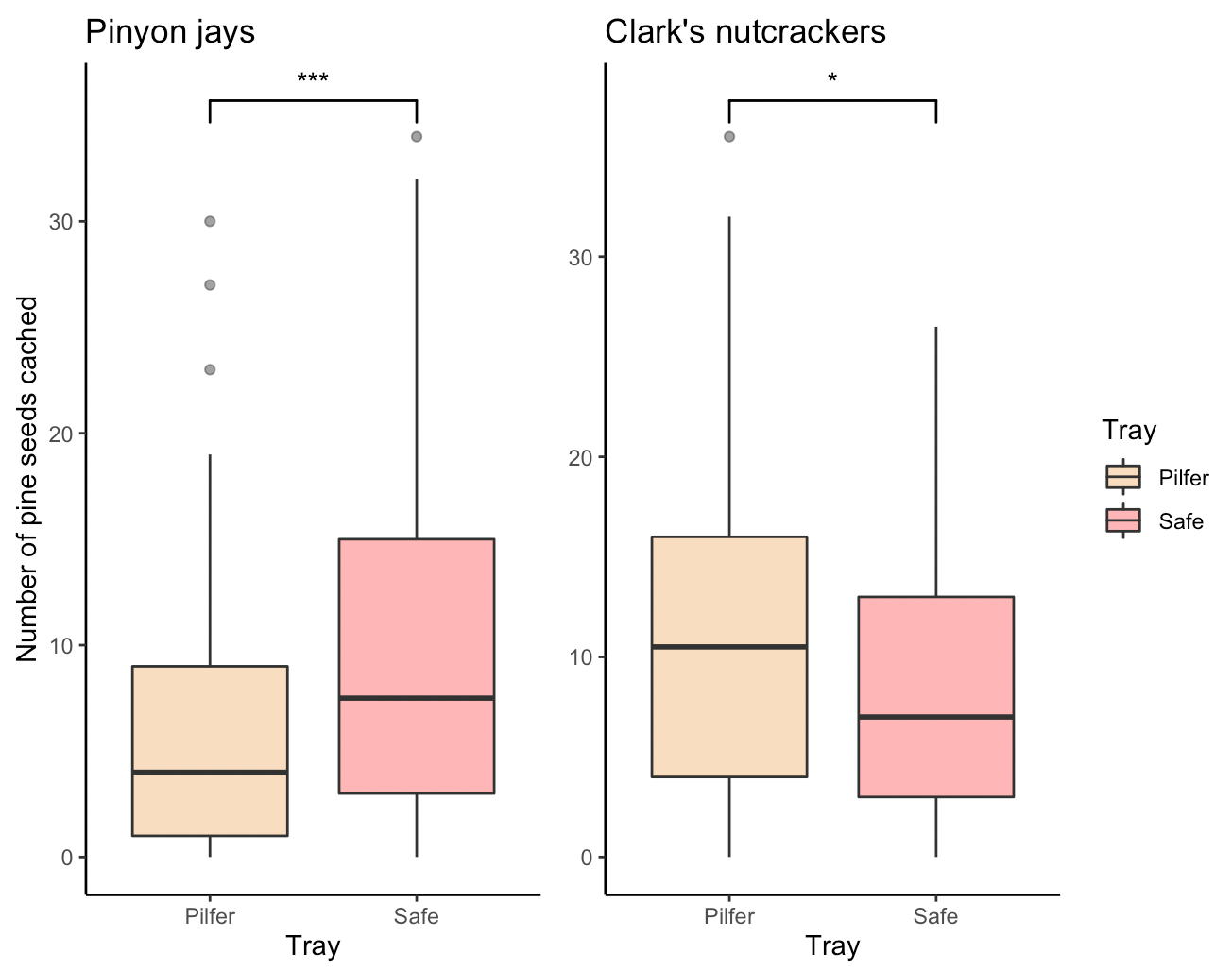

**Figure S2.** Number of pine seeds cached in each tray by pinyon jays (left) and Clark’s nutcrackers (right) during the *Caching phase*. Pinyon jays preferentially allocated their caches in the Safe Tray (*p* < 0.001) and Clark’s nutcrackers preferentially allocated their caches in the Pilfered Tray (*p* = 0.004).

**Figure S3.** Number of pine seeds cached in the Pilfered Tray and in the Safe Tray across blpcks by pinyon jays (left) and Clark’s nutcrackers (right) during the *Caching phase*. There was no statistical support that the number of pine seeds cached in each tray differed between them across blocks for pinyon jays (*p* = 0.334), but there was some for nutcrackers (*p* = 0.010), that was mostly driven by an increase in pine seeds cached in the Pilfered Tray and a decrease in pine seeds cached in the Safe Tray during the last block (*p* < 0.001).

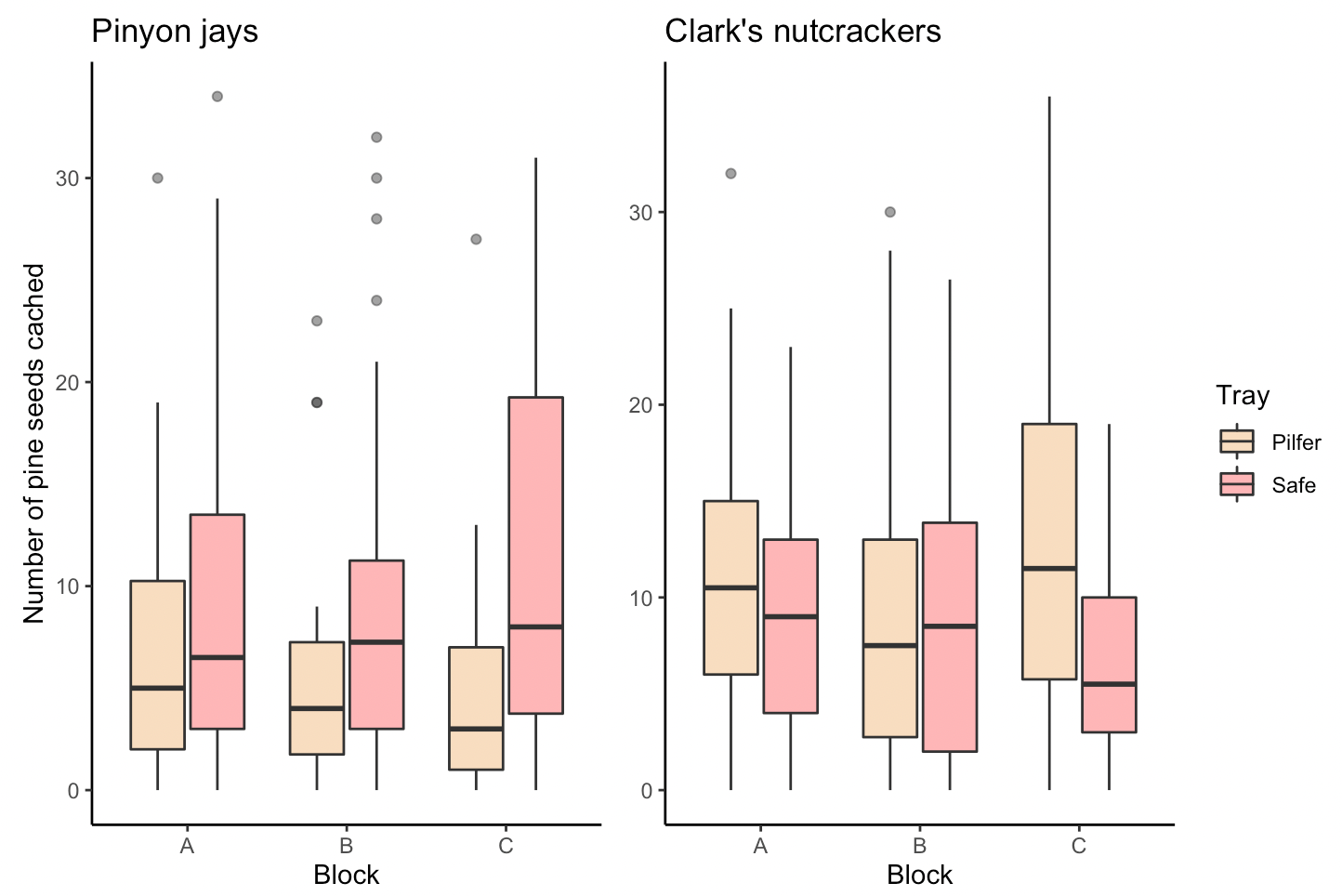

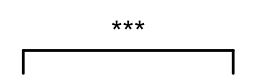

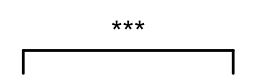

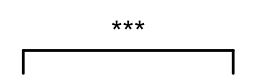

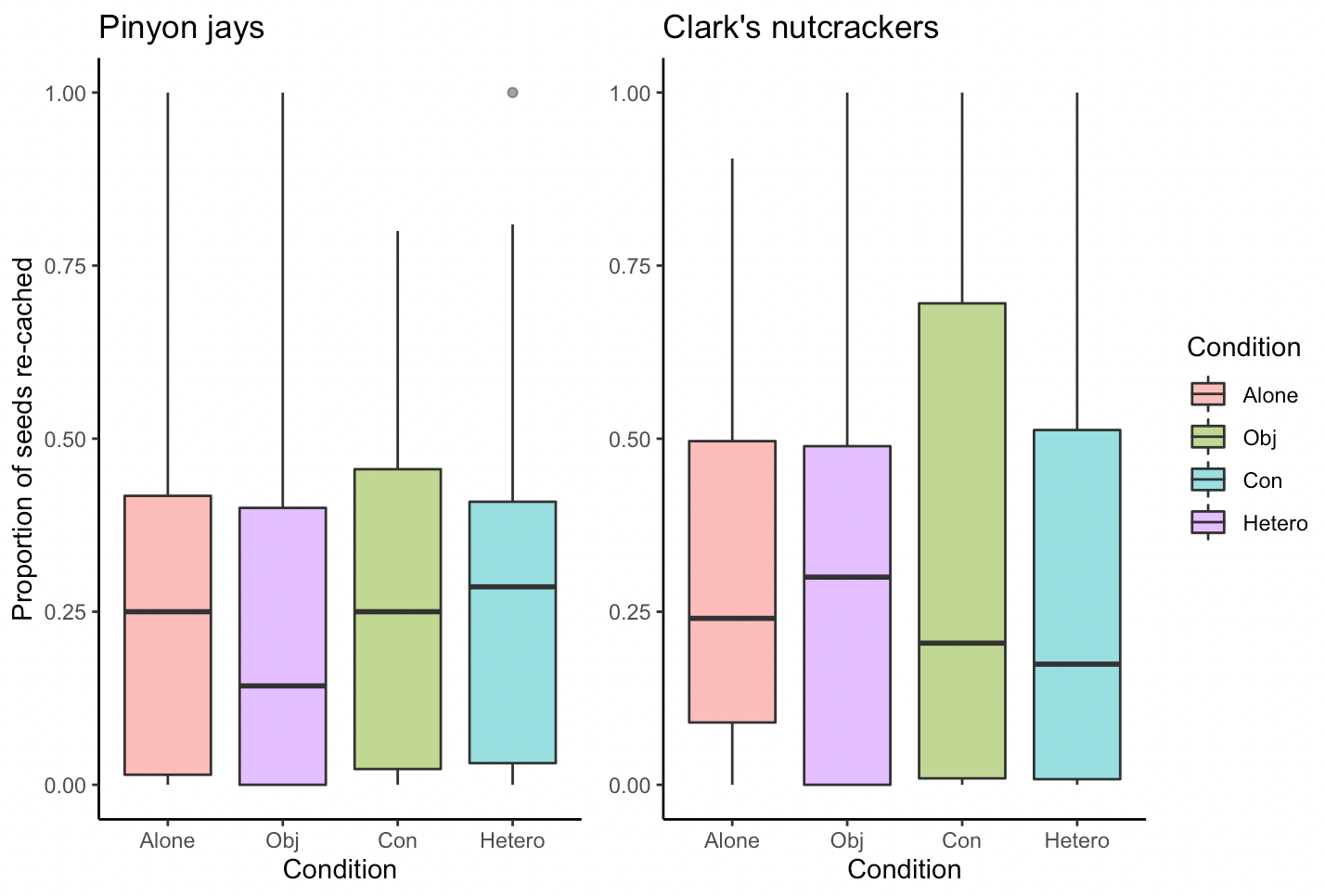

**Figure S4.** Overall proportion of seeds re-cached by pinyon jays (left) and Clark’s nutcrackers (right) during the *Re-caching* *phase*. There was no statistical evidence that the overall proportion of seeds re-cached differed between conditions in pinyon jays (*p* = 0.605) nor in Clark’s nutcrackers (*p* = 0.051).

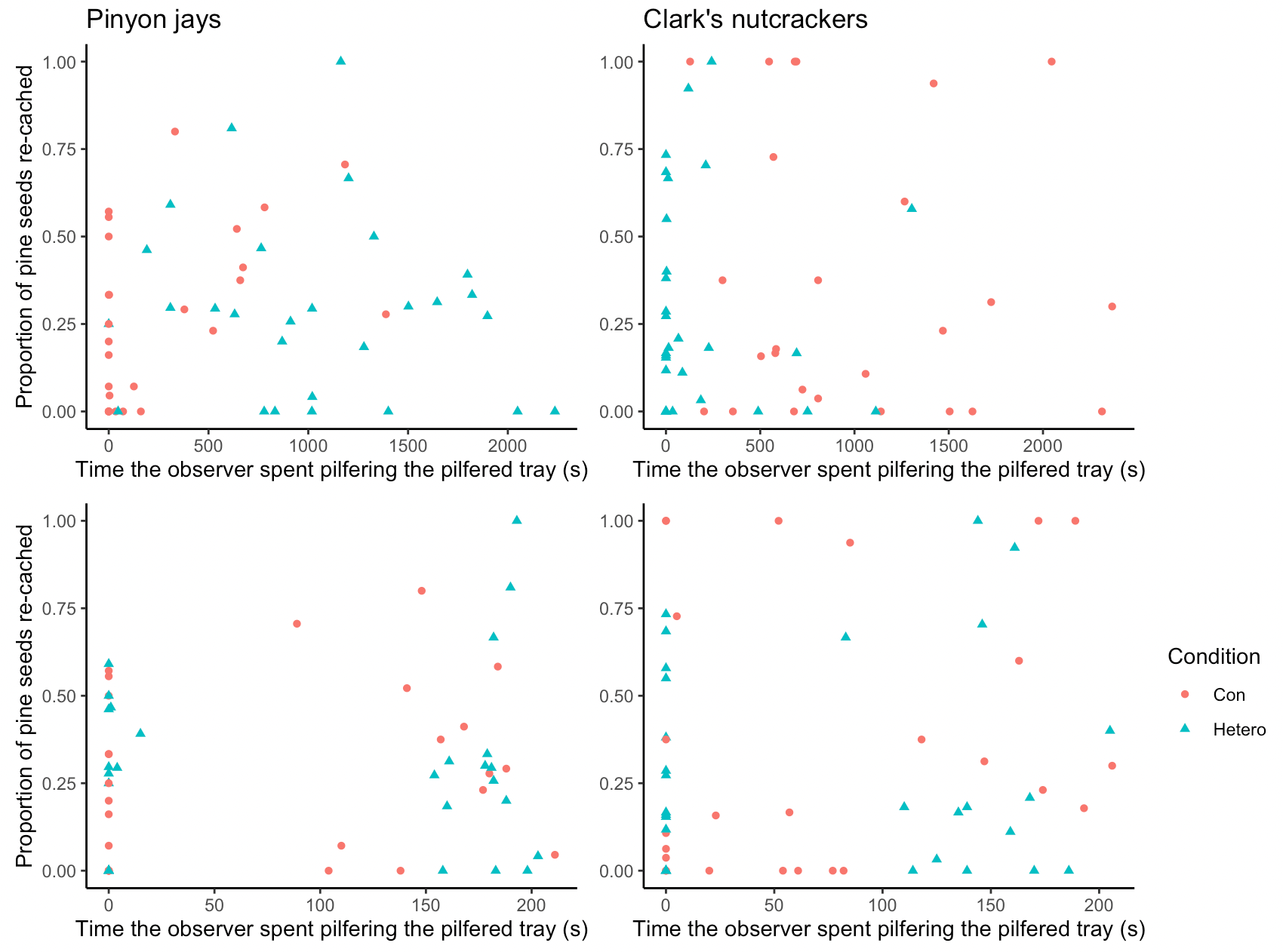

**Figure S5.** Overall proportion of pine seeds re-cached by pinyon jays (left) and Clark’s nutcrackers (right) during the *Re-caching phase* depending on the amount of time the observer spent interacting with the cacher (top) and on the amount of time the observer spent pilfering the Pilfered Tray (bottom). For pinyon jays, the overall proportion of pine seeds re-cached increased with the amount of time the observer spent interacting with the cacher, depending on the condition (*p* < 0.001), but not on the amount of time the observer spent pilfering the pilfered tray (*p* = 0.067). For Clark’s nutcrackers, the overall proportion of pine seeds re-cached increased with the amount of time the observer spent pilfering the pilfered tray, depending on the condition (*p* = 0.005), but not on the amount of time the observer spent interacting with the cacher (*p* = 0.201).

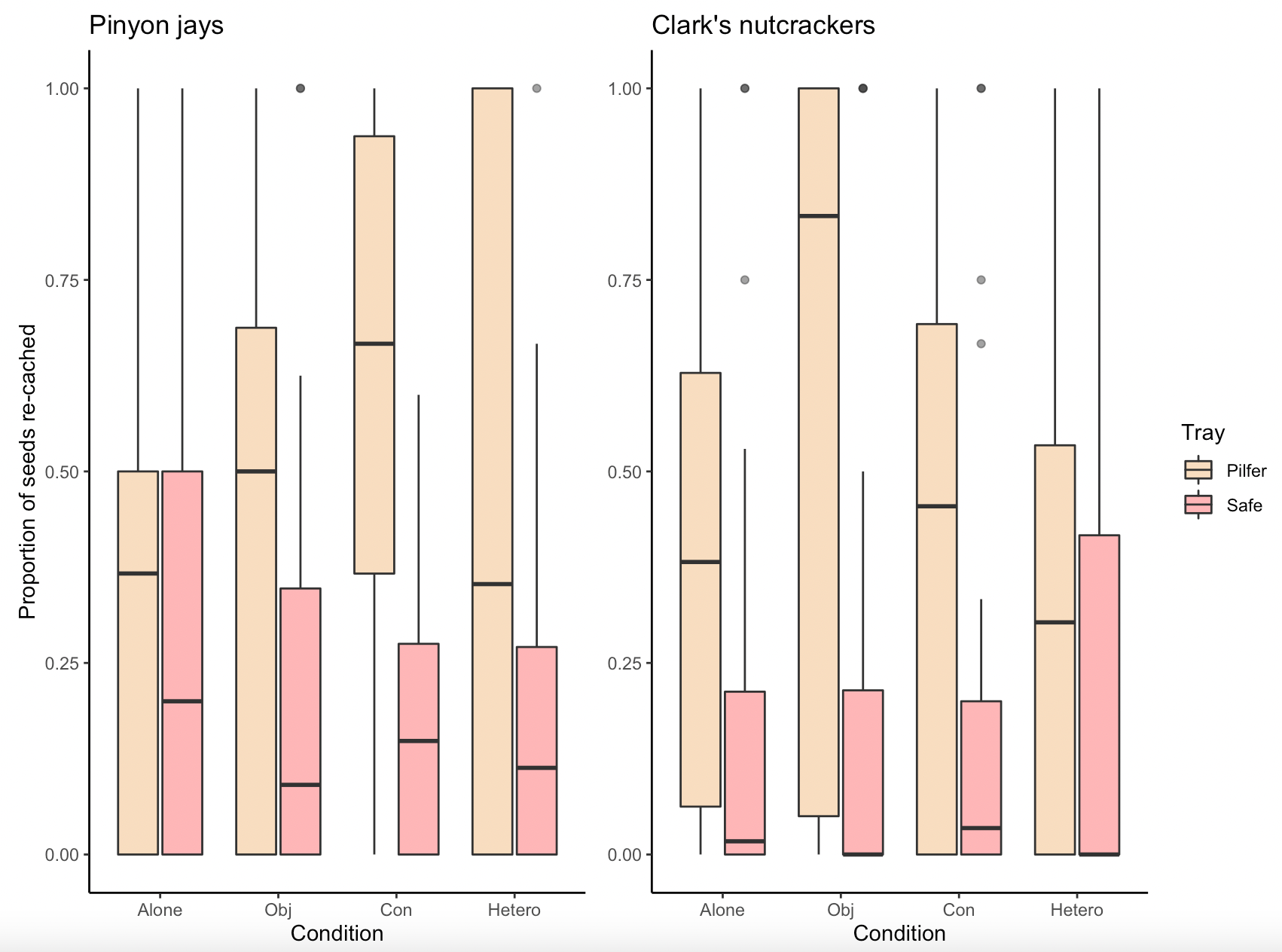

**Figure S6.** Proportion of seeds re-cached in the Pilfered Tray and in the Safe Tray by pinyon jays (left) and Clark’s nutcrackers (right) during the *Re-caching* phase. There was no statistical evidence that the proportion of re-cached seeds per tray differed between conditions in pinyon jays (*p* = 0.811) nor in Clark’s nutcrackers (*p* = 0.083).
